# Tools and Methods from the Anopheles 16 Genome Project

**DOI:** 10.1101/011205

**Authors:** Aaron Steele, Michael C. Fontaine, Andres Martin, Scott J. Emrich

## Abstract

The dramatic reduction in sequencing costs has resulted in many initiatives to sequence certain organisms and populations. These initiatives aim to not only sequence and assemble genomes but also to perform a more broader analysis of the population structure. As part of the Anopheline Genome Consortium, which has a vested interest in studying anpopheline mosquitoes, we developed novel methods and tools to further the communities goals. We provide a brief description of these methods and tools as well as assess the contributions that each offers to the broader study of comparative genomics.

## 1 Introduction

In recent years the cost of sequencing genomic data has decreased from nearly $10,000 per megabase (Mb) in 2001 to less than $0.10 per megabase today. This dramatic reduction in cost has spurred many initiatives to sequence certain organisms or groups of organisms at great depth [3] [10] [12]. With this wealth of new data, novel analysis techniques and tools are constantly being developed to extract useful information.

As part of the Anopheline Genome Consortium (AGC) [15] we developed several tools and methods for Next Generation Sequencing (NGS) data. To encourage adoption of our methods and to streamline analysis of others, we present our methods and underlying code as a publicly available toolkit (https://bitbucket.org/steelea/16genometoolkit) available for use and redistribution under the GNU Public License. More detailed documentation of the methods and tools is provided at https://bitbucket.org/steelea/16genometoolkit/wiki/Home.

## 2 Methods

### The Pipeline

For the AGC consortium we developed a variant discovery pipeline based heavily on the GATK’s best practices[4, 7, 14]. Similar to GATK, Our pipeline is divided into 4 major steps and is designed to discover high quality variants even with relatively low coverage samples. The four major step of the pipeline are Alignment, Pre-processing, Variant Calling, and Variant Recalibration. We provide a brief description of each stage below and direct the reader to https://bitbucket.org/steelea/16genometoolkit/wiki/Pipeline for the full list of commands and parameters contained therein.

1. **Alignment** - The alignment step aligns paired-end short reads to a reference assembly using BWA’s[13] aln algorithm. The alignments are cleaned by updating the MAQ0 for soft-clipped reads. The cleaned reads are then sorted by their reference coordinate using Picard Tools[2].
2. **Pre-processing** - The pre-processing phase performs additional scrubbing of the aligned reads. Optical duplicates are marked and read group information is added. In this step we also perform local realignment of reads to insure optimal mapping of reads near insertions and deletions. At this step GATK’s best practices suggest that Base Quality Score Recalibration (BQSR) should also be performed, however, given the nature of low coverage samples and a lack of biologically verified variants, we found it best to omit this step to avoid skewing downstream analysis.
3. **Variant Calling** - For variant calling we recommend using either the HaplotypeCaller of UnifiedGenotyper packaged within GATK. Either can be used with a reasonable amount of confidence, but there are definite runtime and quality trade-offs between the two. Even though GATK is recommended for variant calling any tool that produces variants in the VCF format can be substituted without issue.
4. **Variant Recalibration** - The variant recalibration step is the most important step and is where we deviate the most from GATK’s best practices. To recalibrate and find high quality variants we first annotate the variants for several metrics (i.e. read depth, inbreeding-coefficient, allele balance, etc.). Hard filters are applied to the variants to retain only biallelic SNPs that have a reasonable quality and depth and are found in a given percentage of the samples. These variants deemed to be of the highest quality (most confident) are then used for the Gaussian Mixture Model Recalibration step as outlined in GATK’s best practices.

## 3 Tools

In this section we describe the major tools developed for various analysis. Because new tools are continuously being developed, we classified them into the following four major categories: Data Acquisition, Variants, Phylogeny, and Population Genomics.

### 3.1 Data Acquisition

#### sraDownloader

NCBI recently incorporated the Aspera Connect[1] plugin into its website to simplify the process of acquiring data from SRA. Aspera Connect is browser plugin that makes the process of getting data from SRA as simple as clicking a “download” link. Unfortunately, this makes the process of acquiring data in a terminal (Unix/Linux) environment painstakingly difficult. With the Aspera Connect plugin, you are not only able to download in a terminal environment, but you also no longer have the ability to view an FTP link for simple copy paste download. To overcome these issues we developed the sraDownloader tool which allows for simple downloading of SRA data in a Unix/Linux environment. The primary input is a an SRA accession number of a CSV file that details experiment accession numbers and group information. Using this information the sraDownloader attaches to SRA via FTP to acquire the desired samples without the use of a browser or the Aspera Connect plugin.

### 3.2 Variants

All of the tools in this section are dedicated to the production and manipulation of VCF[3] files.

#### fastaToVCF

The fastaToVCF tool takes a haploid FASTA file and creates a diploid VCF file where each site matches the reference allele. This tool allows the genome assembly of an out-group to be converted to VCF format without the inconvenience of sequencing and variant calling. An example of where this tools is useful is the ABBA test of introgression [6], which requires variants for an out-group. If the out-group variants do not exist this tool allows for the creation of artificial out-group variants from an out-group assembly.

#### vcfToFasta

The vcfToFasta tool takes a diploid VCF file and creates a haploid FASTA file, selecting the majority allele at each site. This tool incorporates variation information into FASTA files that can ultimately be used for multiple sequence alignment or phylogeny construction. This tools is most useful when NGS data is available for several closely related organisms, only one of which has a genome assembly. By calling variants for each species and creating its representative FASTA, which can cautiously be used with genome comparison tools to infer a phylogeny.

#### vcfIdize

The vcfIdize is a tool that adds variant IDs to each variant with a VCF file. Despite the triviality of the script performing the Variant Recalibration step of the pipeline is impossible without it. Additionally, many existing tools that filter variants require that each variant has a unique ID to filter properly.

#### pipe-job-builder

The pipeline introduced in Section 2 is presented in 4 steps, but actually has more than a dozen different intermediate steps to run. To simplify the process of using the pipeline we created the pipe-job-builder script which automatically builds SGE jobs that execute the presented pipeline. The pipe-job-builder simplifies the process of generating jobs for multiples samples as easy as running a single command, but requires a very precise directory structure. Details on the directory structure and the execution of each step in the pipeline are available in the online documentation.

#### hc-builder

There has been extensive discussion of the runtime and quality trade-offs between the HaplotypeCaller and the UnifiedGenoptyper of GATK. In general the HaplotypeCaller tends to produce better variants but the runtime can be restrictive for lots of samples or high-depth samples. Therefore, it is recommended that if you have a sizable set of sequence data you should use the UnifiedGenotyper, despite the lower quality variants it produces. To remedy this we created the hc-builder, which improves the runtime of the Haplotype caller by launching parallel disjoint SGE jobs. The level of parallelization that can be achieved is directly related to the number of contigs/scaffolds in the reference genome. However, even a modest speed-up of 2-4x can be substantial when the projected serial runtime is on the order of days.

### 3.3 Phylogeny

#### mtTree

mtTree is a self-contained miniature pipeline that constructs a phylogeny based on mitochondrial (mtDNA) or ribosomal DNA (rDNA). The pipeline is loosely based on [16], but incorporates more robust tools and specialized scripts for achieving a more accurate phylogeny. Figure 1 shows the workflow of the mtTree pipeline.

**Figure 1:**
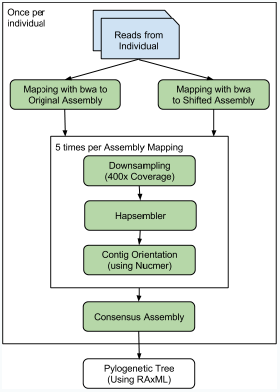
mtTree Pipeline

For each sample short-reads are aligned to a mtDNA or rDNA assembly using BWA[13]. mtTree creates an alternate “shifted” assembly by connecting the two ends of the assembly and breaking it in the middle. The short-reads are then aligned again to the “shifted” assembly to overcome lowcoverage areas that occur at the breakpoints of circular genomes. The reads that aligned to either assembly are extracted and randomly down-sampled multiple times. From each down-sampling a new assembly is constructed using Hapsembler[8]. Each assembly is aligned to the original assembly using Nucmer[11] and a consensus sequence for each sample is generated. These consensus sequences are used in conjunction with RAxML to construct a phylogeny.

### 3.4 Pool-Seq

#### ldx-wrapper

LDx[9] is a tool for estimating linkage disequilibrium (LD) from pooled sequencing data. LDx has already been incorporated into many studies, but the serial nature of the algorithm can introduce a prohibitive runtime overhead, especially for large fragmented reference genomes. To circumvent these runtime limitations we developed the ldx-wrapper script which uses multi-threading to parallelize the running of LDx. In the ldx-wrapper the reference is split based on contigs/scaffolds and each piece is processed by LDx separately using multiple threads.

#### estimMerger

Pool-hmm[5] is a tool for estimating allele frequencies(AF) and detecting selective sweeps in pooled sequencing data. Pool-hmm was designed to estimate allele frequency for a single pool, but offers little in the way of comparing multiple pooled samples. For comparison of different pools we developed the estimMerger which takes multiple .estim files produced by Pool-hmm and merges them into a single tabular file. The resulting output promotes direct comparison of allele frequency between different pools and allows for easy incorporation into other AF-based analyses.

## 4 Conclusions

We present a pipeline for variant discover using short-read data and a collection of bioinformatics tools to aid in comparative genomic studies. The tools are all publicly available for use, improvement, and redistribution. The tools and methods described herein can be acquired from https://bitbucket.org/steelea/16genometoolkit.

## References

[1] Aspera connect plugin. http://downloads.asperasoft.com/connect2/.

[2] Picard tools. http://picard.sourceforge.net/index.shtml. Accessed: 2014-08-30.

[3] GR Abecasis, D. Altshuler, A. Auton, LD Brooks, RM Durbin, R. A. Gibbs, M.E. Hurles, G.A. McVean, DR Bentley, A. Chakravarti, et al. A map of human genome variation from population-scale sequencing. Nature, 467(7319):1061–1073, 2010.

[4] G.A. Auwera, M.O. Carneiro, C. Hartl, R. Poplin, G. del Angel, A. Levy-Moonshine, T. Jordan, K. Shakir, D. Roazen, J. Thibault, et al. From fastq data to high-confidence variant calls: The genome analysis toolkit best practices pipeline. Current Protocols in Bioinformatics, pages 11–10, 2013.

[5] S. Boitard, R. Kofler, P. Françoise, D. Robelin, C. Schlötterer, and A. Futschik. Pool-hmm: a python program for estimating the allele frequency spectrum and detecting selective sweeps from next generation sequencing of pooled samples. Molecular ecology resources, 13(2):337–340, 2013.

[6] Heliconius Genome Consortium et al. Butterfly genome reveals promiscuous exchange of mimicry adaptations among species. Nature, 2012.

[7] M.A. DePristo, E. Banks, R. Poplin, K.V. Garimella, J.R. Maguire, C. Hartl, A.A. Philip-pakis, G. del Angel, M.A. Rivas, M. Hanna, et al. A framework for variation discovery and genotyping using next-generation dna sequencing data. Nature genetics, 43(5):491–498, 2011.

[8] N. Donmez and M. Brudno. Hapsembler: an assembler for highly polymorphic genomes. In Research in Computational Molecular Biology, pages 38–52. Springer, 2011.

[9] A.F. Feder, D.A. Petrov, and A.O. Bergland. Ldx: estimation of linkage disequilibrium from high-throughput pooled resequencing data. PloS one, 7(11):e48588, 2012.

[10] D. Haussler, S.J. O’Brien, O.A. Ryder, F.K. Barker, M. Clamp, A.J. Crawford, R. Hanner, O. Hanotte, W.E. Johnson, J.A. McGuire, et al. Genome 10k: a proposal to obtain whole-genome sequence for 10,000 vertebrate species. The Journal of heredity, 100(6):659–674, 2008.

[11] S. Kurtz, A. Phillippy, A.L. Delcher, M. Smoot, M. Shumway, C. Antonescu, and S.L. Salzberg. Versatile and open software for comparing large genomes. Genome Biology, 5:R12, 2004.

[12] R. Levine. i5k: The 5,000 insect genome project. American Entomologist, 57(2):110–113, 2011.

[13] H. Li and R. Durbin. Fast and accurate short read alignment with burrows–wheeler transform. Bioinformatics, 25(14):1754–1760, 2009.

[14] A. McKenna, M. Hanna, E. Banks, A. Sivachenko, K. Cibulskis, A. Kernytsky, K. Garimella, D. Altshuler, S. Gabriel, M. Daly, et al. The genome analysis toolkit: a mapreduce framework for analyzing next-generation dna sequencing data. Genome research, 20(9):1297–1303, 2010.

[15] D.E. Neafsey, G.K. Christophides, F.H. Collins, S.J. Emrich, M.C. Fontaine, W. Gelbart, M.W. Hahn, P.I. Howell, F.C. Kafatos, D. Lawson, et al. The evolution of the anopheles 16 genomes project. G3: Genes— Genomes— Genetics, 3(7):1191–1194, 2013.

[16] J. Prado-Martinez, P.H. Sudmant, J.M. Kidd, H. Li, J.L. Kelley, B. Lorente-Galdos, K.R. Veeramah, A.E. Woerner, T.D. OConnor, G. Santpere, et al. Great ape genetic diversity and population history. Nature, 499(7459):471–475, 2013.

